# TLR4 competence and mouse models of leptospirosis

**DOI:** 10.1101/2025.02.03.636333

**Authors:** Olifan Zewdie Abil, Suman Kundu, Leonardo Moura Midon, Maria Gomes-Solecki

## Abstract

Mice are slowly being accepted as alternative models for investigation of leptospiral infection. The strain most used to analyse sublethal disease (C3H-HeJ) expresses a *tlr4* gene in its immune cells that is hyporesponsive to leptospiral LPS and thus the model is deemed immunocompromised. To help resolve this valid scientific concern we did a study in which we compared infection with *Leptospira interrogans* serovar Copenhageni Fiocruz in mice expressing fully competent *tlr4* (C3H-HeN, C57BL6) versus *tlr4* hyporesponsive mice (C3H- HeJ) over a period of two weeks. We found that the C3H-HeJ mouse was the only strain that sustained lethal infection, thus confirming the association between *tlr4* function and acute disease. We found that the two mouse strains with a fully functional *tlr4* gene (C3H-HeN and C57BL/6) also developed clinical and molecular signs of sublethal leptospirosis nearly on par with C3H-HeJ, as quantified by weight loss, survival curves, presence of *Leptospira* 16S rRNA in blood and urine, and burden of viable spirochetes in kidney. Regarding kidney markers of inflammation and fibrosis, C3H-HeN and C57BL/6 produced less IL-1 beta, iNOS and ColA1 than C3H-HeJ, which is consistent with increased resilience to infection expected of the *tlr4* competent strains. Regarding Leptospira-specific antibody, all mouse strains produced IgM and T-cell independent IgG3. Interestingly, in contrast to C57BL/6, both C3H strains produced circulating IgG1 but only C3H-HeN produced IgG2a to *L. interrogans* two weeks post infection. Thus, TLR4-competent C3H-HeN and C57BL/6 may be more appropriate mouse models of sublethal leptospirosis, whereas C3H-HeJ can be used to further establish a mouse model of lethal leptospirosis. Our data strongly suggests that TLR4 function is important but not sufficient to cause susceptibility to leptospirosis.

**Author summary:** Previous studies by different laboratories have shown C3H-HeJ to be a susceptible mouse strain to sublethal and lethal leptospirosis. The goal of our study was to evaluate if low *tlr4* responsiveness to leptospiral LPS in C3H-HeJ mice is the factor that drives susceptibility to infection with pathogenic *Leptospira*. We did a comparative study using mouse strains immunocompetent and hyporesponsive to *tlr4*. The data shows that although *tlr4* does affect the virulence of leptospirosis (only C3H-HeJ succumbed to infection), the *tlr4* competent strains (C3H-HeN and C57BL/6) also developed measurable sublethal leptospirosis. Thus, competent recognition of leptospiral-LPS by murine TLR4 plays a role but it is not sufficient to determine susceptibility to leptospirosis. Furthermore, the data suggests that T-cell dependent responses could be involved in susceptibility to leptospirosis in the C3H models of infection.

## Introduction

Leptospirosis, an often overlooked but resurging infectious disease caused by a spirochetal bacteria, is a significant global health concern, impacting millions of individuals worldwide. This disease carries a high mortality rate, resulting in approximately 65,000 deaths annually [1]. Furthermore, it poses a serious threat to animals of agricultural importance, leading to substantial economic losses, particularly in tropical and subtropical regions [2]. Although significant efforts have been made to formulate novel vaccination approaches that confer enduring immunity and safeguard against various serovars, our understanding of the specific immune factors contributing to host defense and disease progression remains limited [3].

Leptospirosis research is challenging due to the inconsistent outcomes observed in leptospiral infections involving small animals, large animals and humans [4] and a plethora of high and low virulence *Leptospira* serovars [5–8]. Hamsters and guinea pigs have been widely used as small-animal models of acute disease, as they recapitulate the manifestations of severe disease in humans. Rats have been used as a model for studying the severity of human leptospirosis as they become chronically infected and shed *Leptospira* in urine many months after infection [9, 10]. However, the usage of these animals in research is limited in some parts of the world where the disease is endemic due to stringent animal regulations [4] and the scarcity of accessible reagents for routine experiments.

Mouse models are extensively utilized in biomedical research because they provide a versatile approach due to the accessibility of diverse immunological and genetic tools, as well as a broad range of available reagents [4]. Researchers have studied the outcomes of experimental leptospiral infection using various mouse models, including studies on lethal, sublethal, and chronic leptospirosis manifestations [4]. For instance, C57BL/6 and BALB/c, exhibit increased resilience to acute disease and have the potential to serve as models for persistent infection caused by *Leptospira interrogans* [11, 12]. C3H-HeJ mice infected with *L. interrogans* develop disease that can be easily monitored through measurement of clinical scores. This strain produces valuable models of lethal [13–15] and sublethal disease [16], and has been used for investigating inflammatory signatures of infection and necroptosis, immunology and efficacy of protection studies [17–19]. Toll-like receptor (TLR) 4, the LPS receptor, was shown to play a central role in the control of leptospirosis [20, 21]. One mechanism described how leptospiral-LPS activates murine, but not human, TLR4 in cultured macrophages [22], and it is associated with resistance to infection [23]. C3H-HeJ mice have a single amino acid substitution (aa712, P to H) within the coding region of the *tlr4* gene that makes this molecule hyporesponsive to the atypical *Leptospira* LPS [21, 24]. Of note, humans, also believed to be susceptible to leptospirosis express a TLR4 molecule that does not sense the atypical *Leptospira* LPS [22]. Nevertheless, one consistent criticism regarding the use C3H-HeJ mice is that its hyporesponsive Toll-Like Receptor 4 (TLR4) qualifies these mice as immunocompromised. To help resolve this valid scientific concern we did a study in which we compared infection with *L. interrogans* in mice expressing fully competent *tlr4* (C57BL6, C3H-HeN) versus mice that are *tlr4* hyporesponsive (C3H-HeJ).

## Material and methods

### Animals

Male, 9-10 week old C3H-HeJ, C3H-HeN, C57BL/6 mice and 7-8 week old C3H-HeJ were used for this study. C3H-HeJ animals were purchased from The Jackson Laboratory (Bar Harbor); C3H-HeN and C57BL/6 were purchased from Charles River (Wilmington, MA) and maintained in specific pathogen-free at the Laboratory Animal Care Unit of the University of Tennessee Health Science Centre, with unrestricted access to food and water. We used male mice because they are more susceptible to leptospirosis [25]. We used 4 animals per group, and experiments were reproduced once. All animal procedures were carried out in accordance with the University of Tennessee Health Science Centre Institutional Animal Care and Use Committee protocol number 22-0362.

### Bacterial strains and culture

*L. interrogans* serovar Copenhageni strain Fiocruz L1-130 (henceforth LiC), frozen in -80°C, was passaged in hamster. Their kidneys were harvested and cultivated in 4mL of Hornsby-Alt-Nally (HAN) media [26] with 100 μg/mL 5-fluorouracil (MP Chemicals, CA) at 29°C for better growth. Passage 2 was done in Ellinghausen-McCullough-Johnson-Harris (EMJH) medium supplemented with Difco *Leptospira* enrichment EMJH (Becton, MD) at 28-30°C. EMJH culture passage 2 was allowed to reach the log phase of growth, pelleted by centrifugation at 3,000 × *g* for 5 min, and washed and resuspended in sterile 1× phosphate-buffered saline (PBS) (Thermo Fisher Scientific). Next, the cells were used to infect the animals after counted under a dark-field microscope (Zeiss USA, Hawthorne, NY) using a Petroff- Hausser chamber as previously described [25].

### Animal infection and collection of specimens

Mice were inoculated with 1×10^8^ LiC in 200 μL of endotoxin-free 1× PBS intraperitoneally (i.p.). Control mice were inoculated with the same volume of endotoxin-free 1× PBS. Survival and body weight loss were monitored for 15 consecutive days post-infection. Mice were euthanized 15 days after post-infection or when they reached the endpoint criteria (20% body weight loss post-infection). Blood, urine, and kidney were collected for further analysis.

### Bacterial quantification

The NucleoSpin tissue kit (Clontech, Mountain View, CA) was used to purify genomic DNA from kidney, blood and urine following the manufacturer’s protocol, and the purified DNA was then stored at −20°C for further analysis. To quantify *Leptospira*, quantitative polymerase chain reaction (qPCR) was performed using *Leptospira* 16S rRNA primers (Forward: CCCGCGTCCGATTAG and Reverse: TCCATTGTGGCCGAACAC) and a TAMRA probe (CTCACCAAGGCGACGATCGGTAGC) obtained from Eurofins (Huntsville, AL). The results were reported as the number of *Leptospira* genome equivalents. The qPCR mixture consisted of 25 μM of each primer, 250 nM of the specific probe, and 2 μL of DNA sample, with a total volume of 20 μL. Duplicate reactions were performed. The amplification protocol included an initial step of 10 min at 95°C, followed by 40 cycles of amplification (15 s at 95°C and 1 min at 60°C). The analysis was conducted using the comparative threshold cycle (CT) method. A negative result was determined if no amplification occurred.

### Measurement of *Leptospira*-specific antibody

*Leptospira*-specific antibody levels were measured using the enzyme-linked immunosorbent assay (ELISA). The process of preparing a leptospiral extract for LiC was carried out following previous instructions [27]. Briefly, LiC was cultured in EMJH media until it reached confluency. The bacterial cells were then separated by centrifugation to form a pellet. This pellet was subjected to incubation with BugBuster® solution (1mL) at room temperature in a shaker (100 rpm) for 20 min and mixed thoroughly by vortexing. The resulting mixture was stored at -20°C. The whole cell extract of *Leptospira* was appropriately diluted in a sodium carbonate coating buffer with a concentration of 1X. A 96-well flat-bottom ELISA microtiter plate (Nunc-eBioscience) was coated overnight at 4°C with 100 µL 1X sodium carbonate coating buffer whole-cell extract of *Leptospira* (10^7^–10^8^ bacteria per well).

After overnight incubation, the ELISA plate was washed with 1X PBST. The plate was blocked by a blocking buffer (100 µL/well) containing 1% BSA, followed by incubation for 1 h at 37°C. After washing, serum samples (1:100) was added, and the plate was incubated for 1 h at 37°C. The unbound primary antibody was removed by vigorous washing. Next, anti-mouse secondary antibodies for IgM IgG, IgG3, IgG1 or IgG2a (all from Cell signaling technology, CST) conjugated with horseradish peroxidase was added, which was incubated for 30 min, followed by standard color development using TMB Sureblue. Absorbance measurement was carried out at OD 450 nm using Molecular Devices Spetramax.

### Expression of inflammatory and fibrosis markers

Total RNA was extracted using the RNeasy Mini Kit (QIAGEN) according to the manufacturer’s protocol. Complementary DNA (cDNA) was synthesized from the extracted RNA using the RevertAid First Strand cDNA Synthesis Kit (Thermo Fisher Scientific). The resulting first-strand cDNA served as the template for reverse transcription PCR (RT-qPCR), which was performed on a QuantStudio™ 3 Real- Time PCR (Applied Biosystems) using the PowerTrack™ SYBR™ Green Master Mix (Applied Biosystems). Each RT-qPCR reaction (10 μL total volume) included the cDNA template and specific primers. The cycling conditions were as follows: an initial step at 50°C for 2 minutes, followed by denaturation at 95°C for 2 minutes, followed by 40 cycles of 95°C for 15 sec for denaturation, and 60°C for 1 min for annealing/extension. A melt curve analysis was conducted at the end of the amplification to confirm the specificity of the PCR products. Relative gene expression levels across samples were quantified by the double delta Ct (2^-ΔΔCt^) method, with glyceraldehyde 3-phosphate dehydrogenase (GAPDH) serving as the endogenous reference control. The primer sequences used in this study are listed in Table S1 of the supplemental material.

### Statistical analysis

We used GraphPad Prism 10 software. The *P-*value was determined by Mann-Whitney *U* test, or Two-way ANOVA with Tukey’s multiple comparison test.

## Results

### Weight loss, burden in blood, shedding in urine and survival after infection with *L. interrogans*

Following experimental leptospiral infection (10^8^ LiC), mice were monitored over a 15-day period (Fig 1A). In C3H-HeJ, a steady weight loss was observed beginning on day 6, reaching a peak around day 9, at which point one mouse met the endpoint criteria. The remaining mice recovered gradually thereafter. The overall weight loss in this group ranged from 2% to 21% (Fig 1D). C3H-HeN mice exhibited daily weight loss from day 1 which decreased gradually until day 14. The overall weight loss in this group ranged from 7% to 16% (Fig 1E). In contrast, C57BL/6 mice experienced low % of weight loss, with an overall decrease of 1% to 8%. (Fig 1F). The weight loss curves measured for the three strains were statistically significant (*p*=0.0116, *p*<0.0001) as compared to their respective control groups (Fig 1D to F).

**Figure 1.**
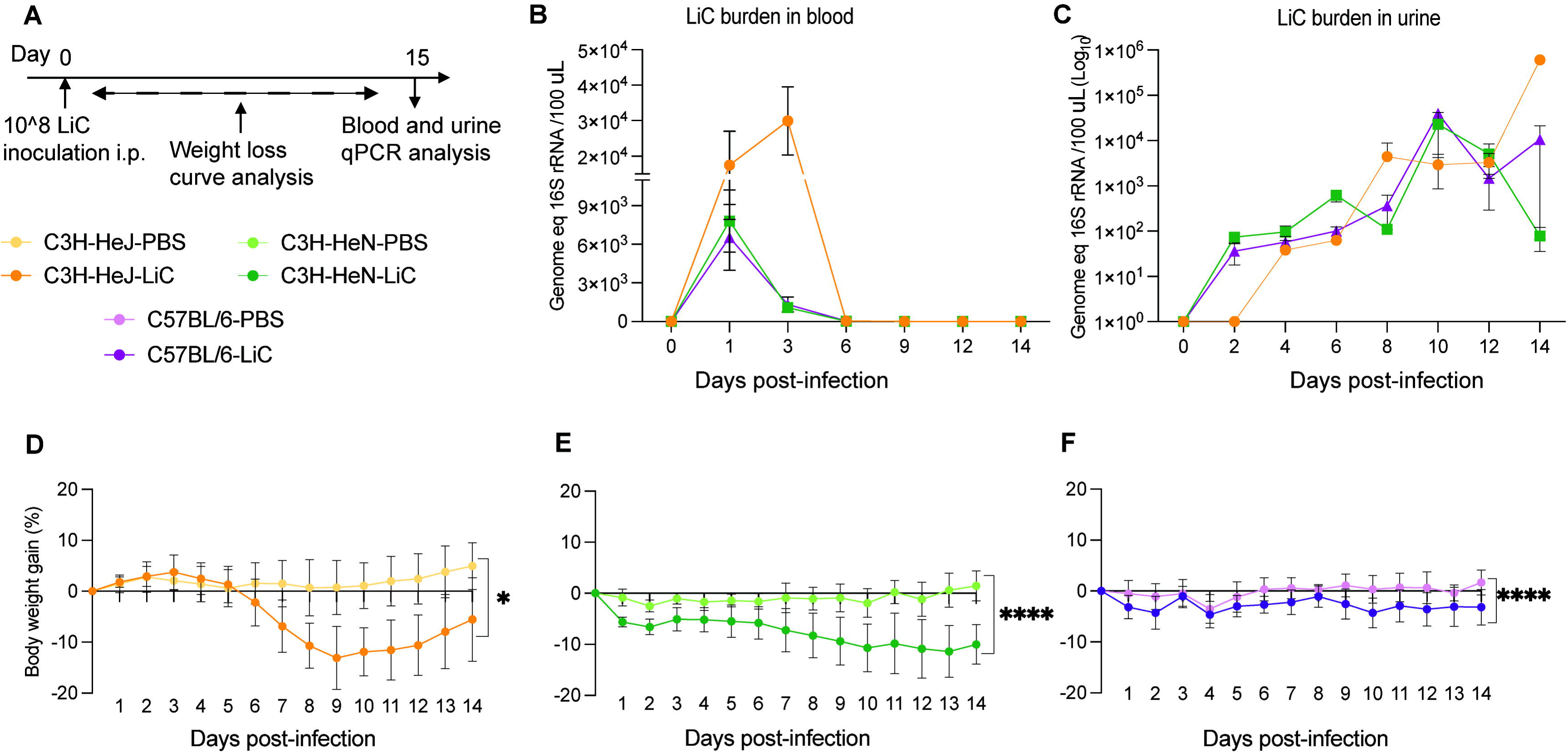
Weightloss and bacterial burden in blood and urine after infection. C3H-HeJ, C3H-HeN and C57BL/6 male mice (n=4/group) were inoculated IP with 1 x 10^8^ *L. interrogans* serovar Copenhageni strain Fiocruz L1-130 (LiC) and with PBS (con­ trol) (A). Quantification of bacterial load in blood (B) and urine (C) was determined by real-time qPCR of the16S rRNA gene. Body weight (^0^/o change) was recorded daily for 15 days after infection (D to F). * p=0.0116,*\*\*\*\*p* < 0.0001 by Mann-Whitney *U* test. Data shown represents one of two experiments.

Quantitative PCR (qPCR) targeting the *Leptospira* 16S rRNA gene was performed to analyse the dissemination of *Leptospira* in blood and urine. Blood samples were collected on days 1, 3, 6, 9, 12, and 14 post-inoculation to quantify the *Leptospira* burden. Peak *Leptospira* burdens were observed on day 3 in C3H-HeJ mice (∼3x10^4^), whereas the peak burden occurred on day 1 in both C3H-HeN and C57BL/6 mice, ∼6-9x10^3^ (Fig 1B). *Leptospira* shedding in urine was also assessed using qPCR on alternate days over a 15-day period. *Leptospira* shedding was detected a low levels (∼10^2^) during the first week post-infection, peaking on day 10 in the second week for C3H-HeN and C57BL/6 mice (∼10^4^), whereas in C3H-HeJ mice it peaked on day 14 at ∼10^6^ (Fig 1C). Regarding survival, the data shows that 75% of C3H-HeJ mice survived at 15 days post-infection, whereas 100% of C3H-HeN and C57BL/6 mice survived during this period (Fig 2).

**Figure 2.**
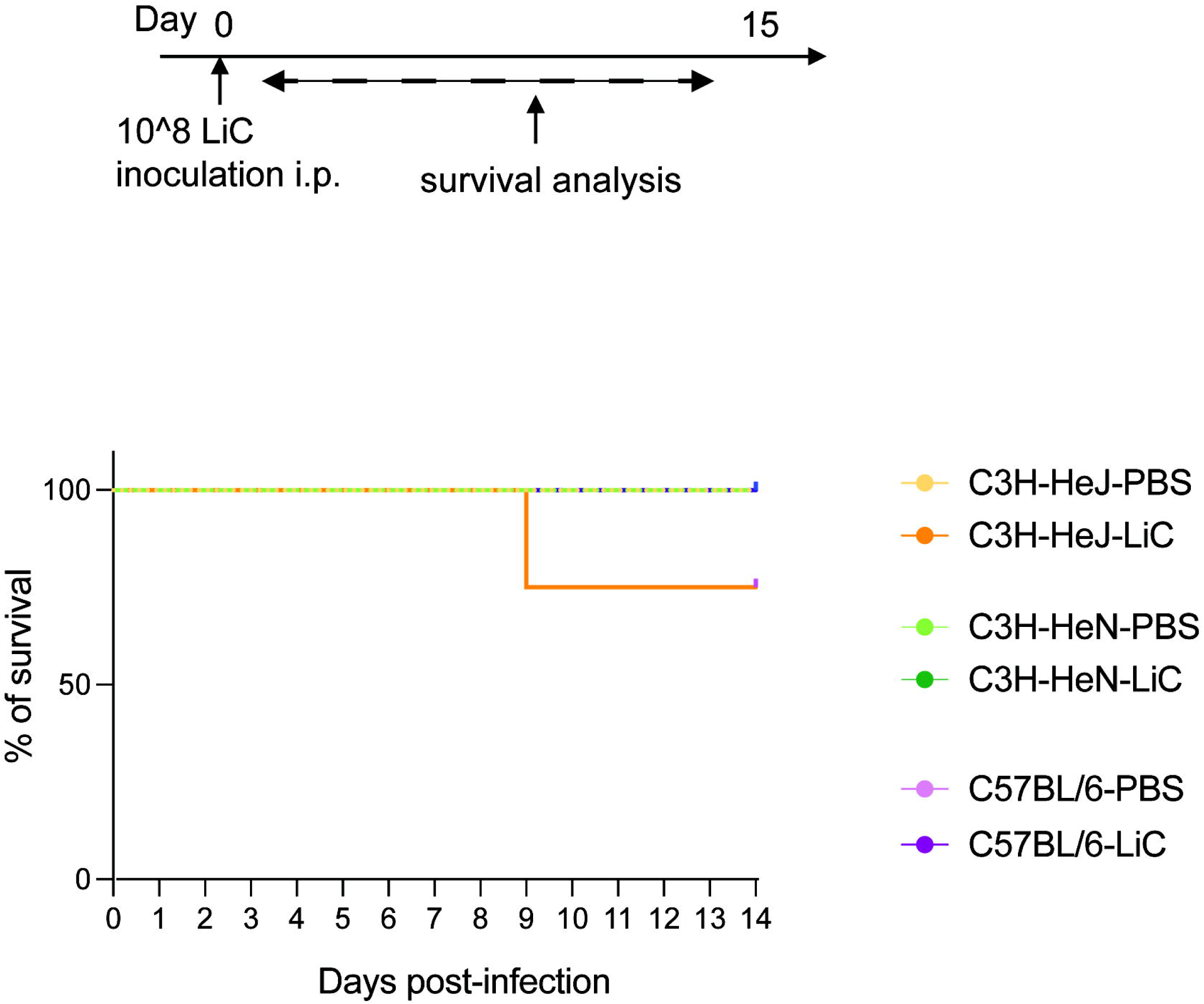
Survival of C3H-HeJ, C3H-HeN and C57BL/6 after infection. Male mice (n=4/group) were inoculated IP with 1 x 1QA8 L *interrogans* serovar Copenhageni Fiocruz L1-130 (LiC) and PBS (control), and the percentage of mice that survived infection for 15 days was recorded. Data represents one of two experiments.

Survival of C3H-HeJ to lethal infection was further explored by using 7-8 week old mice infected with the same dose of *L. interrogans* serovar Copenhageni FioCruz (10^8^) and weight was monitored for 15 days. We found that 100% of the mice reached 20% of weight loss between days 9-12 post infection and met endpoint criteria for euthanasia (Supplemental Fig 1).

### The burden of *L. interrogans* and viability in kidney tissue after infection

To assess renal colonization by *L. interrogans*, kidney tissues from infected mice were collected on the day the mice were euthanized on day 15 post-infection and analyzed for the presence of *Leptospira* DNA using qPCR (Fig 3A). The bacterial burden was ∼7.5 × 10⁴ *Leptospira* per mg of kidney tissue in C3H-HeJ, ∼4.1 × 10⁴ in C3H-HeN, and ∼2.1 × 10⁴ in C57BL/6 infected mice (Fig 3B). Additionally, the viability of *Leptospira* was assessed by culturing kidney tissues in EMJH medium at 30°C for 4 days and quantifying viable spirochetes using qPCR. On day 4 of culture, the average number of viable spirochetes per 100 μL of EMJH culture was ∼4.3x10^3^ for C3H-HeJ, ∼3.1 × 10³ for C3H-HeN, and ∼2.2 × 10³ for C57BL/6 mice (Fig 3C).

**Figure 3.**
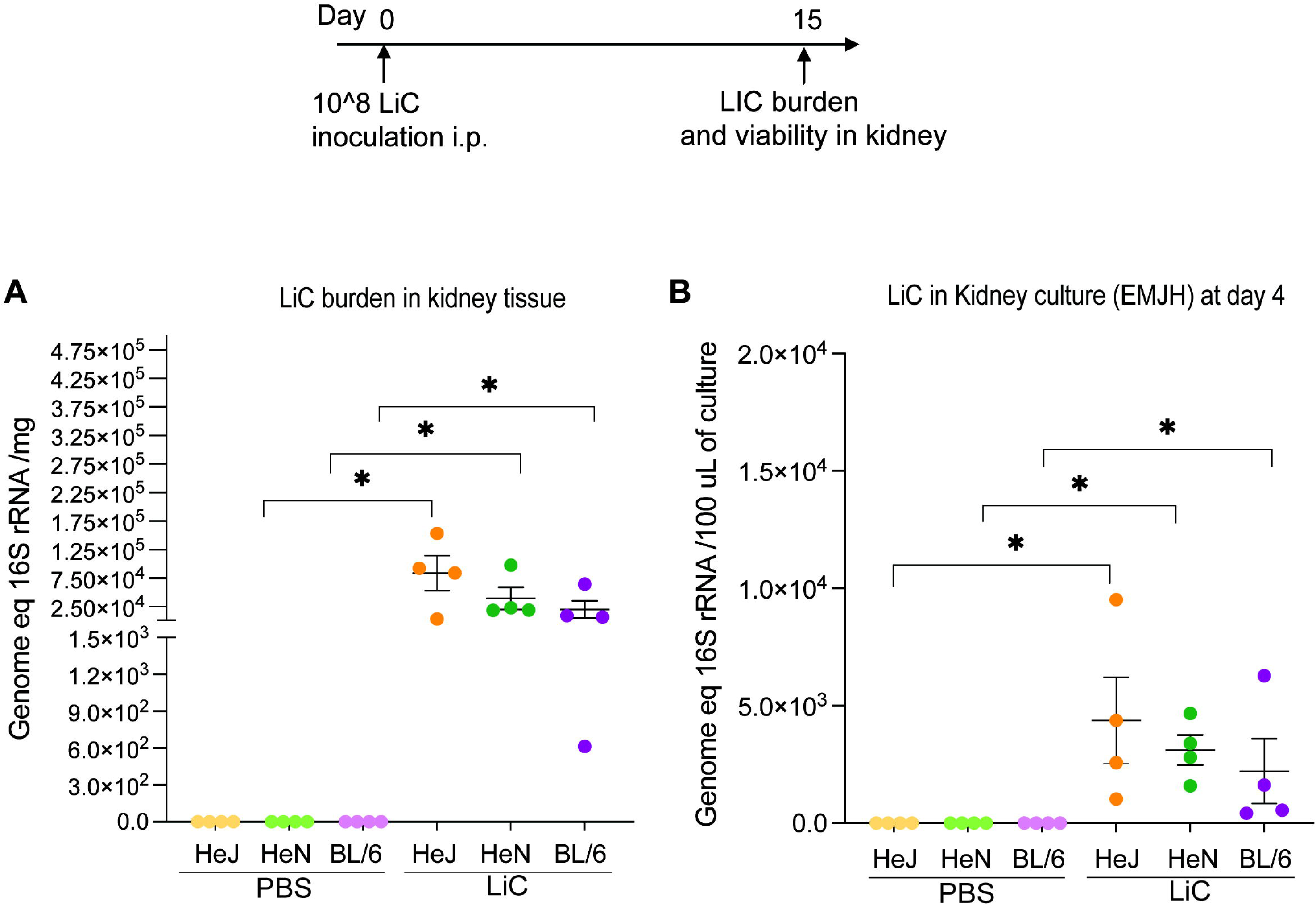
Burden and viability ofL *interrogans* in kidney. C3H-HeJ, C3H-HeN and C57BL/6 male mice (n=4/group) were inoculated IP with 1 x 10“8 L *interrogans* serovar Copenhageni Fiocruz L1-130 (LiC) and with PBS (control). Kidney tissues were collected 2 weeks post infection for qPCR analysis of 16S RNA from DNA purified from kidney tissue (A) and from DNA purified from kidney culture in EMJH (B). Each dot represents a single mouse. *\*p* < 0.05 was determined by Mann-Whitney *U* test. Data represents one of two experiments.

### Expression of inflammatory cytokines and signatures of fibrosis in kidney of C3H-HeJ, C3H-HeN and C57BL/6 mice following *L. interrogans* infection

To assess the upregulation of inflammatory cytokines critical for immune responses and fibrosis markers associated with kidney damage, mice were inoculated with *L. interrogans* as described in Fig 4. Kidney tissues were collected and processed for mRNA expression analysis 15 days post-infection. The expression levels of IL-1 beta mRNA, a key cytokine involved in inflammatory responses was significantly upregulated both in C3H-HeJ and C3H-HeN mice (Fig 4B). Importantly, mRNA of the fibrosis marker, inducible nitric oxide synthase (iNOS), was significantly increased in both C3H-HeJ and C3H-HeN mice, but not in C57BL/6 (Fig 4C). The mRNA expression of collagen A1 (ColA1) was also markedly elevated in both C3H-HeJ and C3H-HeN mice and no ColA1 upregulation was detected in C57BL/6 (Fig 4D). We also analysed the mRNA expression levels of IFN-gamma, IL-17, IL-6, and IL-23 cytokines in all mouse strains; however, the differences observed were not statistically significant.

**Figure 4.**
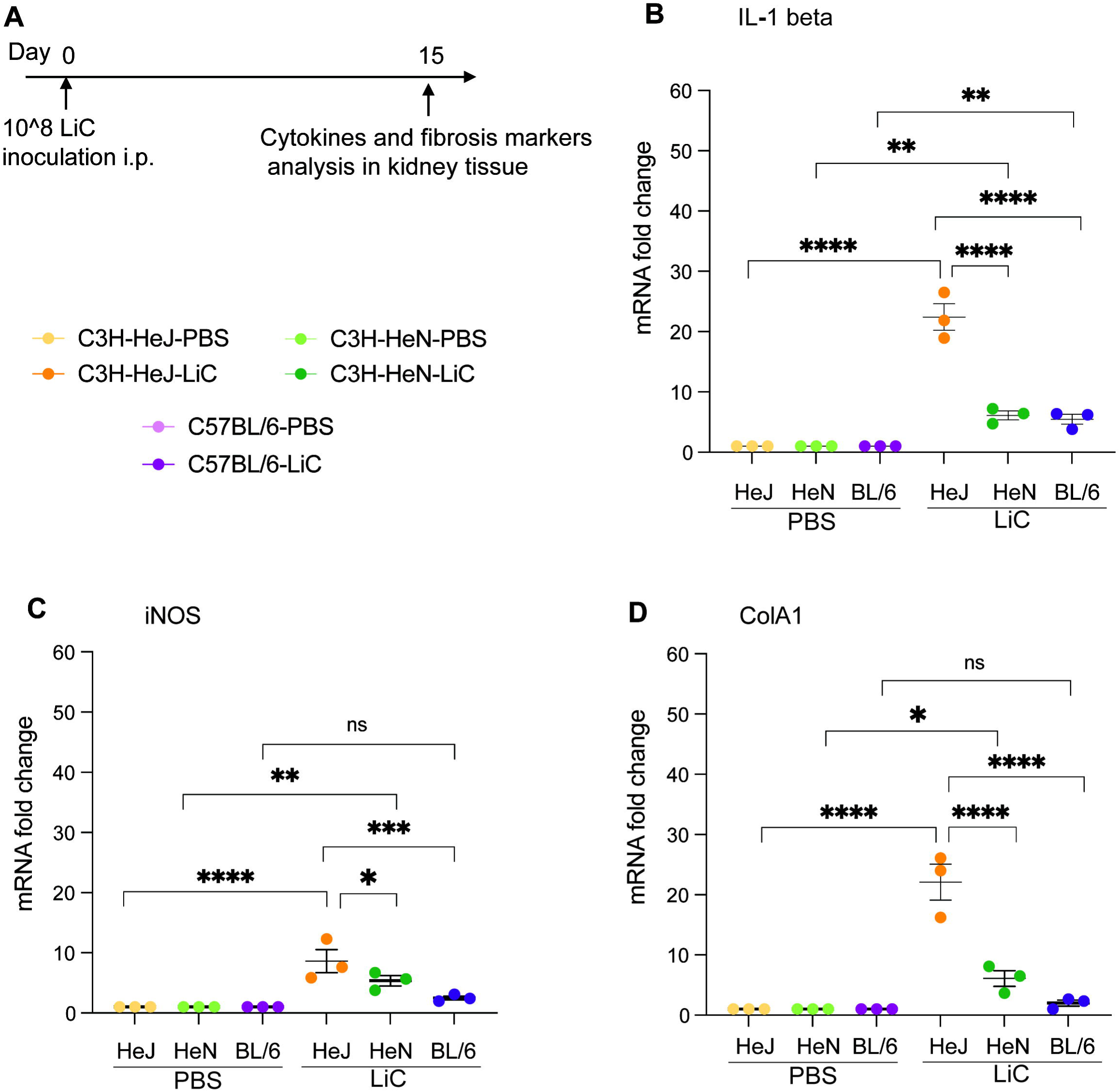
Inflammatory cytokines and fibrosis markers in kidney of infected mice. The expression of IL-1 beta (B), iNOS (C) and ColA1 (D) in kidney tissue were measured using RT-PCR. Glyceraldehyde 3-phosphate dehydrogenase (GAPDH) was used as the calibrator gene to normalize the expression of genes. Each dot represents measurement from a single mouse. *\*p<* 0.05, *\*\*p<* 0.005, *\*\*\*p<0.0005* or *\*\*\*\*p* < 0.0001 was determined by two-way ANOVA followed by Tukey’s multiple comparison correction test. Data shown represents one of two experiments.

### Serological antibody responses in C3H-HeJ, C3H-HeN and C57BL/6 at 15 days post infection with *L. interrogans*

Terminal blood samples were collected 15 days post-infection to evaluate the levels of antibody production to *L. interrogans* (Fig 5). As expected, the three strains of mice produced increased levels of anti-*Leptospira*-specific IgM (Fig 5B), and IgG (Fig 5C) compared to the respective uninfected controls. IgG isotyping produced the following results: IgG3 was the only antibody significantly produced by the three mouse strains (Fig 5D) and IgG1/IgG2a were not detected in C57BL/6 (Fig 5E and F). Significantly higher levels of IgG1 were measured in C3H-HeJ than C3H-HeN compared to the respective controls (Fig 5E) and IgG2a was measured in both C3H strains but was only significant in C3H-HeN (Fig 5F).

**Figure 5.**
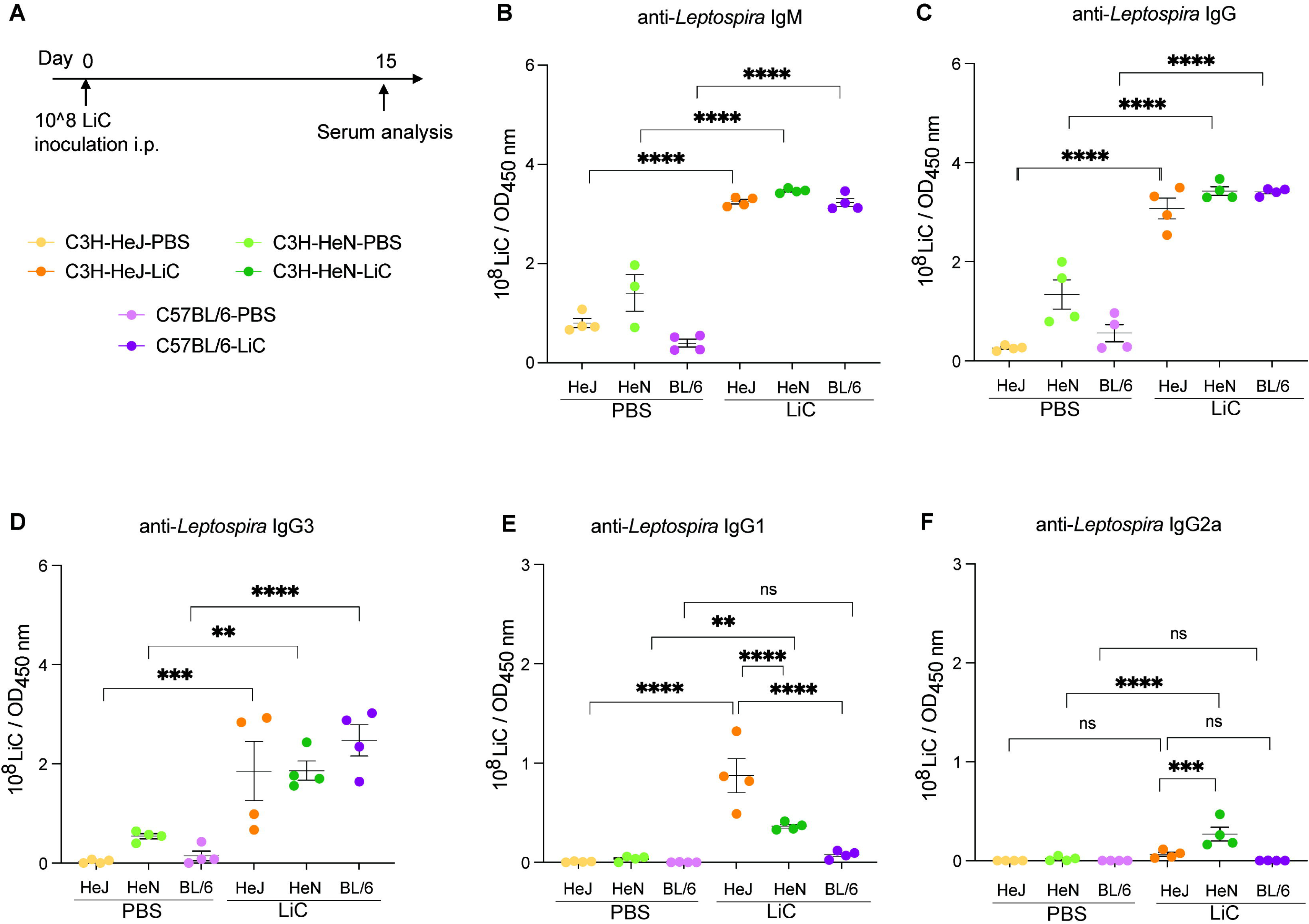
Antibody responses toL *interrogans* infection. C3H-HeJ, C3H-HeN and C57BL/6 male mice (n=4/group) were inoculated IP with 1 x 10^8^ *L. interrogans* serovar Copenhageni Fiocruz L1-130 (LiC) and with PBS (control) (A). Serologic leptospiral-specific lgM (8), lgG (C), lgG3 (D), lgG1 (E) and lgG2a **(F)** was measured using ELISA. Each dot represents a single mouse. *\*p* < 0.05, *\*\*p* < 0.01, *** *p* < 0.001 or *\*\*\*\*p* < 0.0001 was determined by two-way ANOVA followed by Tukey’s multiple comparison correction test. Data shown represents one of two experiments.

## Discussion

We previously used the C3H-HeJ strain to develop a mouse model of experimental sublethal leptospirosis [16, 25, 28]. Since resistance to lethal acute infection with *L. interrogans* was associated with *tlr4* [23] and C3H-HeJ is hyporesponsive to *L. interrogans* atypical LPS, there are valid concerns that this mouse model does not recapitulate a fully competent immune response to this spirochete. In this study, we did a side-by-side comparative analysis using three mouse strains (C3H-HeJ, C3H-HeN and C57BL/6). C3H-HeJ was the only strain that sustained lethal infection thus confirming an association between a fully functional *tlr4* gene and susceptibility to acute *L. interrogans* infection. Nevertheless, the two mouse strains with a fully functional *tlr4* gene (C3H-HeN and C57BL/6) also developed clinical and molecular signs of sublethal leptospirosis over two weeks of infection, nearly on par with C3H-HeJ. Our data indicates that TLR4-competent C3H-HeN and C57BL/6 may be more appropriate mouse models of sublethal leptospirosis, whereas C3H-HeJ may be good model of lethal leptospirosis.

We found weight loss and survival differences between C3H and C57BL/6 in that C3H mice showed larger weight loss curves (∼ 2-21%) than C57BL/6 mice (1-8%) throughout the 15- day infection period, that in the case of C3H-HeJ, affected survival. For a choice of animal models of disease, differences in weight loss in the C3H background produced clinical scores that are easier to reproduce consistently. qPCR analysis of *Leptospira* 16S rRNA demonstrated differences in the dynamics of *L. interrogans* dissemination in blood and urinary shedding among the 3 mouse strains that culminated in a gradient of kidney colonization which was highest in C3H and lowest in C57BL/6. This is consistent with the severity of systemic dissemination observed for C3H strains in weight loss and survival, suggesting that overall, C3H mice are more susceptible to *Leptospira* infection than C57BL/6.

There are numerous links between competent recognition of Leptospiral LPS by TLR4 and resistance to infection in mice [20–23]. Our study confirms that hyporesponsivenes to *tlr4* results in lethal infection in C3H-HeJ mice (Fig 1, Fig 2 and Supplemental Figure 1). Our data also shows that susceptibility to sublethal infection can be observed in C3H-HeN and C57BL/6 mice, which express fully competent TLR4s, although differences in measurements are more readily obtained in C3H-HeN. Of note, humans, also believed to be susceptible to leptospirosis express a TLR4 molecule that does not sense the atypical *Leptospira* LPS [22]. In another study we found that inoculation of potentially susceptible TLR4/MD-2 humanized transgenic mice with *L. interrogans* did not produce different measurements of disease progression or *Leptospira* dissemination than wildtype mice [29]. This may have been related to the humanized transgenic mouse background being C57BL/6. Overall, the data suggests that although TLR4 plays a role in susceptibility to *Leptospira* infection, it is not the ultimate determinant as other receptors in mammalian cells (ex. TLR2) likely compensate for deficient receptor binding.

While innate immune responses are the main immunological pathway for eliminating *Leptospira*, the humoral immune response plays a vital role in effectively eradicating the bacteria and expelling it from the host [30]. Our present observations revealed that LiC infection led to an increase of *Leptospira*-specific IgM, IgG antibodies in the blood of the three mouse strains. This was expected as several other groups have reported [16, 23, 31, 32]. However, IgG isotyping in C3H models of infection produced some interesting, unexpected results for IgG3, IgG1 and IgG2a. IgG3 was the only antibody significantly produced by the three mouse strains and may explain high total IgG for C57BL/6 in our study. Two weeks after infection of C57BL/6, IgG1 was extremely low as observed by others [32]. IgG2a antibodies were not detected in C57BL/6 as can be explained by the absence of a gene for IgG2a [33]. However, at this early time point of infection (2 weeks) significantly higher levels of IgG1 were measured in C3H-HeJ than C3H- HeN, and IgG2a was measured in both C3H strains but was only significantly higher in C3H- HeN. The production of Th2-dependent IgG1 antibody in C3H-HeJ and C3H-HeN mice suggests a Th2-biased activation of the immune response, as shown in our previous studies for C3H-HeJ [16]. Presence of Th1 dependent IgG2a in C3H-HeN mice at two weeks post infection with *L. interrogans* suggests that this response may be driven in part by TLR4 which has been associated with resistance and may explain why disease progression in this mouse is less pronounced than C3H-HeJ. The data strongly suggests that the mouse genetic background affects the type of IgG antibody response to pathogenic *Leptospira*; hyporesponsiveness to TLR4 may account for lower levels of IgG2a in the C3H-HeJ genetic background which result from impaired Th1 responses and could explain increased susceptibility to leptospirosis.

In conclusion, our data suggests C3H-HeN and C57BL/6 mice, both TLR4 competent strains, can be used to recapitulate sublethal leptospirosis, although the C3H-HeN strain produces unambiguous differences in clinical and molecular measurements of disease progression, tissue pathology and bacterial dissemination. Furthermore, and considering that both C3H-HeJ and humans are hyporesponsive to leptospiral LPS, the C3H-HeJ strain may be more suitable for analysis of lethal leptospirosis on par with the hamster model. In addition, both C3H strains can be used to investigate competent Th immune responses to pathogenic *Leptospira* in early time-points of infection. Each of the mouse strains have different characteristics that can be leveraged in pursuit of knowledge on immunity and host response to pathogenic *Leptospira*.

## Acknowledgments

This work was supported by NIAID grant numbers R21 AI142129 (MGS), R44 AI167605 (MGS) and R01 AI175417 (MPI MGS). The content of this manuscript is totally the responsibility of the authors and does not involve the official views of NIH NIAID.

## Author contributions

MGS designed the study. OZA, SK and LMM performed the experiments and data analysis. OZA, SK and MGS wrote the paper.

