## Supplementary figures and images for "TLR4 competence and mouse models of leptospirosis"

### Supplementary Figure 1

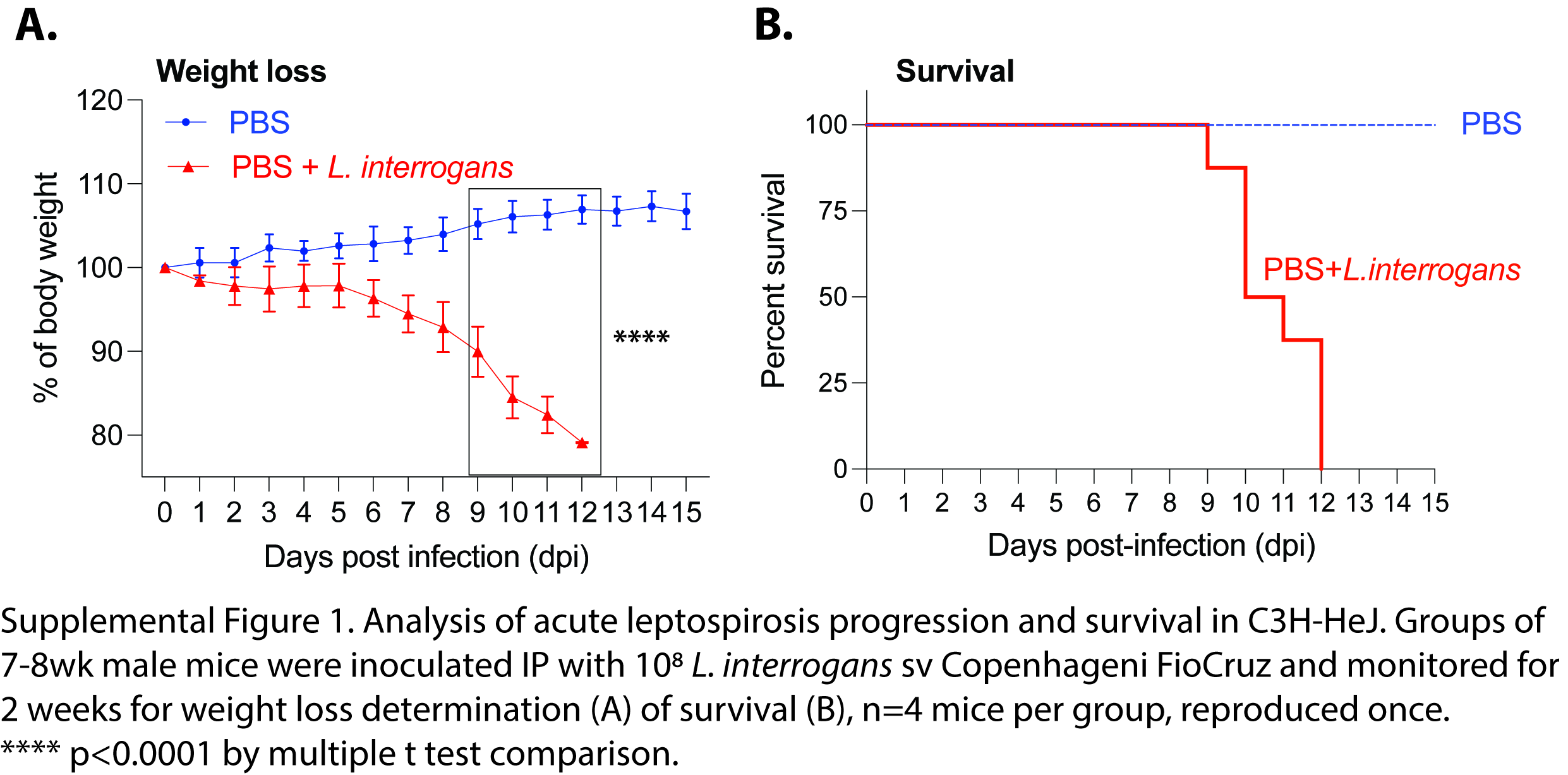
